# Mural-VISTA: a tool for mural cell-vessel interaction assessment and multiscale single-cell topo-morphological analysis

**DOI:** 10.64898/2026.08.27.747487

**Authors:** Hedele Zeng, Mingzhao Hu, Li-Kun Phng, Yukiko T. Matsunaga

## Abstract

Three-dimensional (3D) mural cell morphology is heterogeneous and coupled to vessel geometry, however, measurements from two-dimensional (2D) maximum intensity projections (MIP) obscure overlapping processes and cell-vessel contacts. Accordingly, we developed Mural-VISTA, a semi-automated Python workflow for mural cell-vessel interaction and single-cell topo-morphology analysis of reconstructed surface meshes. This workflow integrates mesh pretreatment, interactive centerline extraction, hierarchical segmentation of cell soma, main axis and secondary processes (branches), and extraction of 37 multiscale (cell process segment level, process level, and whole cell level) topo-morphological and vessel-referenced metrics. Mural-VISTA identified morphological changes in pericytes and vascular smooth muscle cells (vSMCs) with altered RhoA activity. Constitutively active RhoA (RhoA CA) over-expression reduced branch complexity and increased alignment in both cell types, while increased whole-cell and branch solidity only in vSMCs. Dominant-negative RhoA (RhoA DN) over-expression increased branch abundance and reduced branch solidity in pericytes but not vSMCs, suggesting cell-type specific effect of reduced RhoA activity. In conclusion, Mural-VISTA enables quantitative 3D profiling of mural cell architecture and its spatial relationship with the vessel.

## INTRODUCTION

Mural cells, consisting of pericytes (PCs) and vascular smooth muscle cells (vSMCs), associate with the vascular endothelium and contribute to endothelial cell (EC) support, vascular stablization, extracellular matrix (ECM) homeostasis, and regulation of blood flow through contractile activity [1]. PCs are predominantly distributed on capillary beds and exhibit an elongated or branched shape that conform to the vessel surface [2]. Meanwhile, vSMCs typically settle on larger diameter vessels (arteries/arterioles and veins/venules) and form a seamless layer to provide contractile support [1]. These distinct architectures suggest that mural cell morphology is closely linked to its function and associated vascular location.

Advances in 3D in vivo imaging and tissue clearing technology have revealed that mural cells are structurally more diverse than that suggested by conventional 2D imaging. PC populations differ in biomarkers, such as NG2, Desmin and PDGFR-beta, morphology and cell functions [3]. Specifically in brain, PC subtype zonation phenomenon relies on the vessel position and branch level [4]. Calcium dynamic patterns also vary across vascular zones and subtypes to conform to the microenvironment [5]. Meanwhile, vSMCs exhibit phenotypical plasticity between “contractile” and “synthetic” states, accompanied by changes in cytoskeletal organization and cell shape [6]. Some studies specifically point out that mural cell topo-morphology can further vary across developmental stages and organ type [7].

Topo-morphological profiling may therefore offer a structural readout of mural cell states and its response to perturbations. Previous studies have demonstrated that the remodelling of mural cell morphology is associated with its function and highly depends on the local microenvironment [8]. For instance, different vascular geometries and blood flow regimes induce distinct shear stress and transvascular force patterns. Shear stress activates ALK/Smad [9] and Notch pathways [10] in endothelial cells (ECs), and these signals are transmitted to mural cells via Notch-mediated communication [11]. Transvascular force also stretches the cell membrane to open Ca^2+^ channels in mural cell, thus triggering cell constriction [5]. Because mural cells conform to curved vascular surfaces, their phenotype cannot be fully described by cell shape alone. The soma, cellular processes branching pattern, main axis and their connections with associated vessel must also be quantified.

Nevertheless, available analysis methods cannot fully characterize the cellular architecture and cell-vessel geometry. Conventional single-cell morphology quantification often works on 2D images or MIPs of 3D confocal images, which collapse depth information and obscure the complexity of mural cell processes and their interaction with blood vessels. 3D mesh imaging and reconstruction techniques from recent studies provided important descriptive information about mural cell shape and microscale structures [12-14]. However, there is still a lack of a unified tool that aids quantitative and multiscale analysis. Established tools for neurons and microglia, such Simple Neurite tracer [15], ProMoIJ [16], MotiQ [17], and 3DMorph [18], enable process tracking, geometric metrics calculations and allows 3D analysis. However, mural cells have structural anisotropy and conform to associated vessel shape [2]. Existing methods that need manual marking [15] are time-consuming in complex mural cells, in contrast, methods that use global threshold [18] lack vessel-referenced metrics and are unsuitable to mural-cell-specific hierarchical segmentation. Additionally, above morphological skeleton-based methods do not capture surface-derived properties such as solidity, compactness or vessel-referenced properties such as coverage area. A user-friendly analysis framework that integrates centerline topology, surface morphology and vessel-referenced spatial metrics is therefore needed.

Accordingly, we present Mural-VISTA (Mural cell-Vessel Interaction and Single-cell Topo-morphology Analysis), a Python-based workflow for paired 3D surface meshes (a mural cell and its associated vessel). Mural-VISTA combines mesh preprocessing, interactive extraction of process centerlines, hierarchical segmentation of the cell body, primary and secondary processes, multiscale topological and morphological measurements, and quantification of cell–vessel spatial relationships. We validated Mural-VISTA with cross-method, intra-user and inter-user comparisons, then applied it to characterize zebrafish PCs and vSMCs morphological phenotypes arising from altered RhoA activity. Results showed that Mural-VISTA provides a quantitative framework for analyzing 3D mural cell architectures across cell types and molecular perturbations.

## METHODS

**Figure 1.**
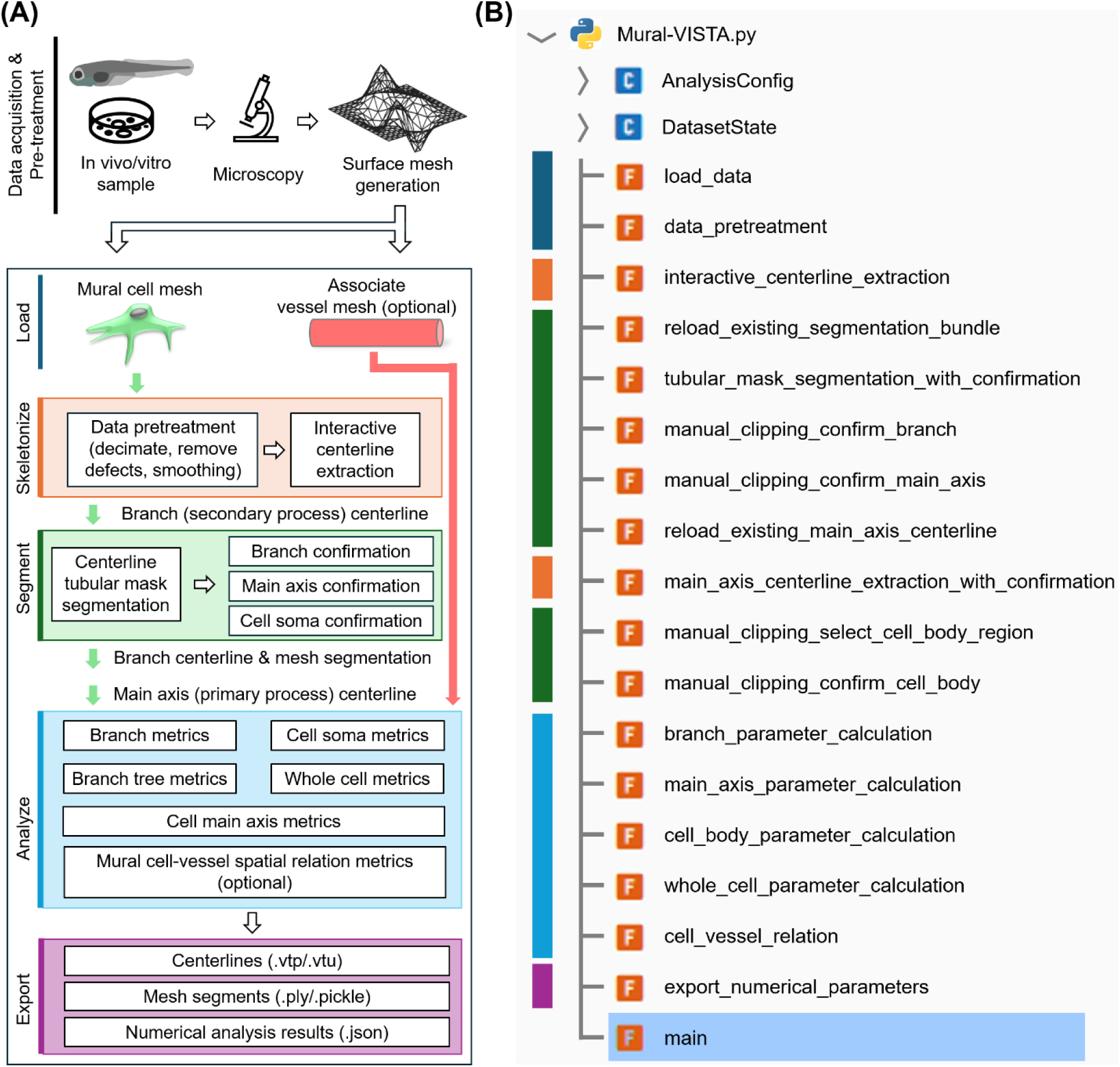
Schematic overview of Mural-VISTA workflow (A) and the corresponding module-level function list (B).

### Software environment

Mesh processing and centerline analysis were performed in a Python 3.10 environment using VTK [19] (v9.2.6), PyVista [20] (v0.45.3), and VMTK [21] (v1.5.0). Mesh defects were repaired with PyMeshFix [22]. Convex hulls were generated by Trimesh (v4.12.2) and alpha shapes were generated using GUDHI [23] (v3.11.0). Graphical user interface (GUI) was generated with PySide6 (v6.8.1) and the Windows executable file was packaged with PyInstaller (v6.21.0) Statistical analysis and calculations were employed using NumPy (v2.2.6) and SciPy (v1.15.2).

### Mesh loading

The user should provide the directory where the mural cell mesh and the associated vessel mesh were stored. The *main* function searches the directory for the .ply files and pairs the mural cell mesh with its associated vessel mesh based on a shared filename prefix. Then, it calls the *load_data*, which will read the paired .ply meshes through a VTK pipeline and check whether the mesh has been processed previously. Unprocessed meshes are decimated before subsequent analysis.

### Mesh pretreatment

The *main* function calls *data_pretreatment*, which opens an interactive GUI for manual mesh fidelity check and trimming. Mesh regions are selected using PyVista *enable_cell_picking* function and the displayed mesh is updated through a customized keyboard-triggered refresh event. A clip can be undone or restarted via internal GUI helper functions *undo_clip* and *refresh_stage* any time before the final confirmation.

After mesh trimming, the *data_pretreatment* function will fix the holes and non-manifold faces using a MeshFix pipe first, calculate the vertex normal, offset a 0.50 µm margin on the normal direction, smooth and subdivide the surface mesh, to guarantee a fine mesh for the robust centerline extraction. The preprocessed mesh is exported as a .ply file with a “_clipped” suffix.

Function *interactive_centerline_extraction* receives the “_clipped” mesh as input and open a GUI to let the user to select the start and the end points of the branches. The GUI inherits from the basic VMTK *vmtkscripts.vmtkCenterlines* interface, but its Python source code was modified to enable customized version to : 1) export the selected start points and end points; 2) keep the start point selection results visible during the end point selection section and 3) differ start point selection label from end point selection using different colors and label sizes.

### Mural cell process centerline extraction

Function *interactive_centerline_extraction* checks the number of points in each extracted centerlines and remove segments only containing one single point. Retained centerlines are resampled at a 0.5 µm interval to standardize centerline point spacing caused by different mural cell mesh size. Then, centerlines are passed through branch extractor, branch merge, and branch geometry calculation filters.

In this study, protrusions were classified into two types as an established study suggested [2]. The primary process, predominantly shaped by apical microtubules and intermediate filaments along the longitudinal direction, was designated as the cell main axis. The secondary processes, predominantly shaped by basal actin filaments along the circumferential direction, were grouped into branch trees. Individual segments within each branch tree were defined as branches.

Branch tree start points were selected at the junctions between the main axis and the branch trees, whereas end points were selected at distal branch tips. The cost function for the fast-marching centerline extraction algorithm was set as 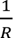 empirically, where *R* denotes the local diameter. The implementation also permits alternative cost functions, such as 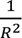 or 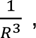 to damp the front propagation speed within some inter-anastomose thinly connected branches, as needed.

Processed meshes bypass function *data_pretreatment* and meshes with readable start/end point lists bypass function *interactive_centerline_extraction*.

### Mural cell mesh segmentation

The *main* function first calls *reload_existing_segmentation_bundle to* check whether a previously generated segmentation bundle is available. If not applicable, *tubular_mask_segmentation_with_confirmation* function coarsely segments the branches from the main axis and the soma. This function uses the centerline trajectory and the maximum inscribed sphere radius on each centerline point to generate an irregular shaped tubular surface VTK Polydata object. Then, it calculates the signed distance distribution from the mural cell mesh to the tubular surface. Vertices enclosed by the tubular surface are assigned negative distances, vice versa. Vertices with a distance below 3 µm are defined as the branch vertices, and the remaining vertices are assigned to main axis and soma region. If the user is unsatisfied with the centerline and the rough segmentation result, this workflow can return to the *interactive_centerline_extraction*. After the automatic tubular segmentation, the user confirms and selects the soma region on main axis regions through functions: *manual_clipping_confirm_branch, manual_clipping_confirm_main_axis, manual_clipping_select_cell_body_region* and *manual_clipping_confirm_cell_body*. These subfunctions open GUIs and enable the user to trim unnecessary regions. Similarly, *reload_existing_main_axis_bundle* attempts to reload previous main axis centerline and switch to call *main_axis_centerline_extraction* if it fails.

### Mural cell mesh morphology analysis

After the segmentation, *branch_parameter_calculation* re-checks the number of points in each segment and deletes single-point centerline. Branch length, branch curvature, branch torsion and branch tortuosity are calculated from the each branch segment. Branch volume and branch surface area are calculated from the tubular surface using the maximum inscribed sphere radius as the local radius.

The branch tree is defined as the connected branches and is identified from the connectivity map. Branch tree max level is defined as the highest branch level within one branch tree. Branch tree fractal dimension is estimated by a box-counting dimension estimation [24]. Branch tree aspect ratio is defined as largest ratio between the oriented bounding box edge lengths. Conceptually related branch tree anisotropy is calculated through a principal component analysis (PCA) where the eigenvalues are defined as *λ*_1_ ≥ *λ*_2_ ≥ *λ*. The anisotropy index is defined as 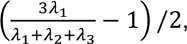 which quantifies the dominance of the first principal component, following established asphericity-based anisotropy shape studies [25,26]. Branch tree max length is defined as the longest path within the tree, while rooted max length denotes the longest path originating from the root. Both are calculated through Dijkstra searching [27]. Conceptually related branch tree sinuosity is defined as the Euclidean distance between the start and end point of the longest path, divided by the longest skeleton path length (branch tree max length). Under this definition, values approaching 1 indicate a locally straighter path. Branch tree straightness is defined as the product of branch tree anisotropy and branch tree sinuosity. Thus, a high straightness value indicates that the branch tree is both globally directional, as reflected by high anisotropy, and locally straight, as reflected by high sinuosity (Table 1).

**Table 1.** Conceptual interpretation of branch tree shape parameters: anisotropy, sinuosity and straightness.

|  | Anisotropy (A) | Sinuosity (S) | Straightness (A×S) | Meaning |
| --- | --- | --- | --- | --- |
| Globally and locally straight | 1 | 1 | 1 | Needle-like branch tree |
| Globally directional but locally curly | High | Low | Medium/Low | Multi-curly-branched branch tree |
| Globally bushy but locally straight | Low | High | Medium/Low | Vessel-wrapping branch tree |
| Globally and locally curly | Low | Low | Very low | Vessel-wrapping, multi-curly-branched branch tree |

Function *main_axis_parameter_calculation* processes the extracted main axis in the same manner as the branch centerlines. Main axis length, curvature, torsion and tortuosity are calculated and exported from this function.

The mesh of soma, or cell body, is sealed as a watertight object and processed by *cell_body_parameter_calculation* and its subfunctions. Soma volume *V* and surface area *A* are calculated directly from the mesh. Soma sphericity is defined as 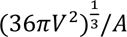 and equivalent diameter is defined as 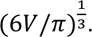 A convex hull will be created to wrap the whole mesh. Soma solidity is defined as ratio that the soma volume divided by the convex volume. Principal axes projection length is defined as the maximum coordinate ranges of the soma mesh vertices along the corresponding PCA axes. Given that soma morphology can provide a structural readout of cellular state [28], we summarized literature-informed interpretation for possible shapes (Table 2).

**Table 2.** Literature-informed interpretation of soma sphericity and solidity.

|  | Nearly spherical soma | Spindle-shaped soma | Concave peanut-shaped soma | Irregular soma |
| --- | --- | --- | --- | --- |
| Sphericity ($\Psi$) | High | Low | Medium | Low |
| Solidity (S) | High | High | Low | Low |
| Interpretation | Pathological or activation [29,30] | Common in homeostatic cells (Baseline) | Undergoing remodeling or cell migration [2] | Pathological or activation [31] |

The whole cell morphological metrics are calculated by *whole_cell_parameter_calculation.* Whole cell volume and surface area are calculated directly from the mesh. Cell solidity is defined as the ratio between mural cell surface mesh volume and its convex hull volume. Cell branch solidity is defined as the ratio between mural cell surface mesh volume and its alpha shape volume. Alpha value is the distance threshold below which the closest 5% of randomly selected vertex pairs fall. The created alpha shape is supposed to cover the intervals between branches while following the overall cell shape. Figure S3 demonstrated the difference between the two types of solidities. Whole cell compactness has a similar definition 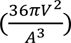 compared with sphericity, but without a cube root operation to amplify the difference and decrease risk of variable confusion. Whole cell Euler characteristic is calculated as the mesh point number minus the edge number and add the face number. Whole cell integral absolute mean curvature (IAMC) is defined as the sum of the absolute mean curvature at each vertex weighted by its associated surface area, following a previous study suggested [32]. Whole cell Willmore energy is similarly defined as the integral area-weighted squared mean curvature on vertices, as another previous study proposed [33]. Both IAMC and Willmore energy have been used in biological, especially for cell membrane and tissue sheet, shape analysis [34-36]. They characterize the complexity of the mural cell structure.

### Mural cell-vessel spatial relation analysis

Function *branch_vessel_relation* quantifies the branch-vessel spatial relationships and uses the vessel mesh as a reference for measuring mural cell and branch orientation. A one-thousand vertices point cloud is randomly subsampled from the reconstructed vessel mesh and fitted to a quadratic Bézier curve. The branch-centerline angle of a branch is defined as the mean angles between each branch element vector and the vessel centerline curve at the nearest point on that curve.

Function *whole_cell_vessel_relation* quantifies the mural cell-vessel spatial relationships. The signed distance between the mural cell mesh and the vessel mesh is calculated same as the segmentation section.

The coverage region is defined by signed distances below -0.3 µm, and the corresponding surface area is reported as the cell–vessel coverage area. The projection length of the whole mural cell on the vessel central curve is also calculated.

### Result exportation

Function *export_numerical_parameters* exports all the numerical analysis results in branch level, branch tree level, whole cell level, as .json files in a subfolder named by the current file name. Extracted centerlines and mesh segments were stored in .vtp and .ply formats for visualization, respectively. They are also stored as Python .pickle packages for faster program reprocessing. The .pickle packages are stored in the same folder with a file name prefix. In total, 37 types of parameters are available and can be exported by this program (Figure 2).

**Figure 2.**
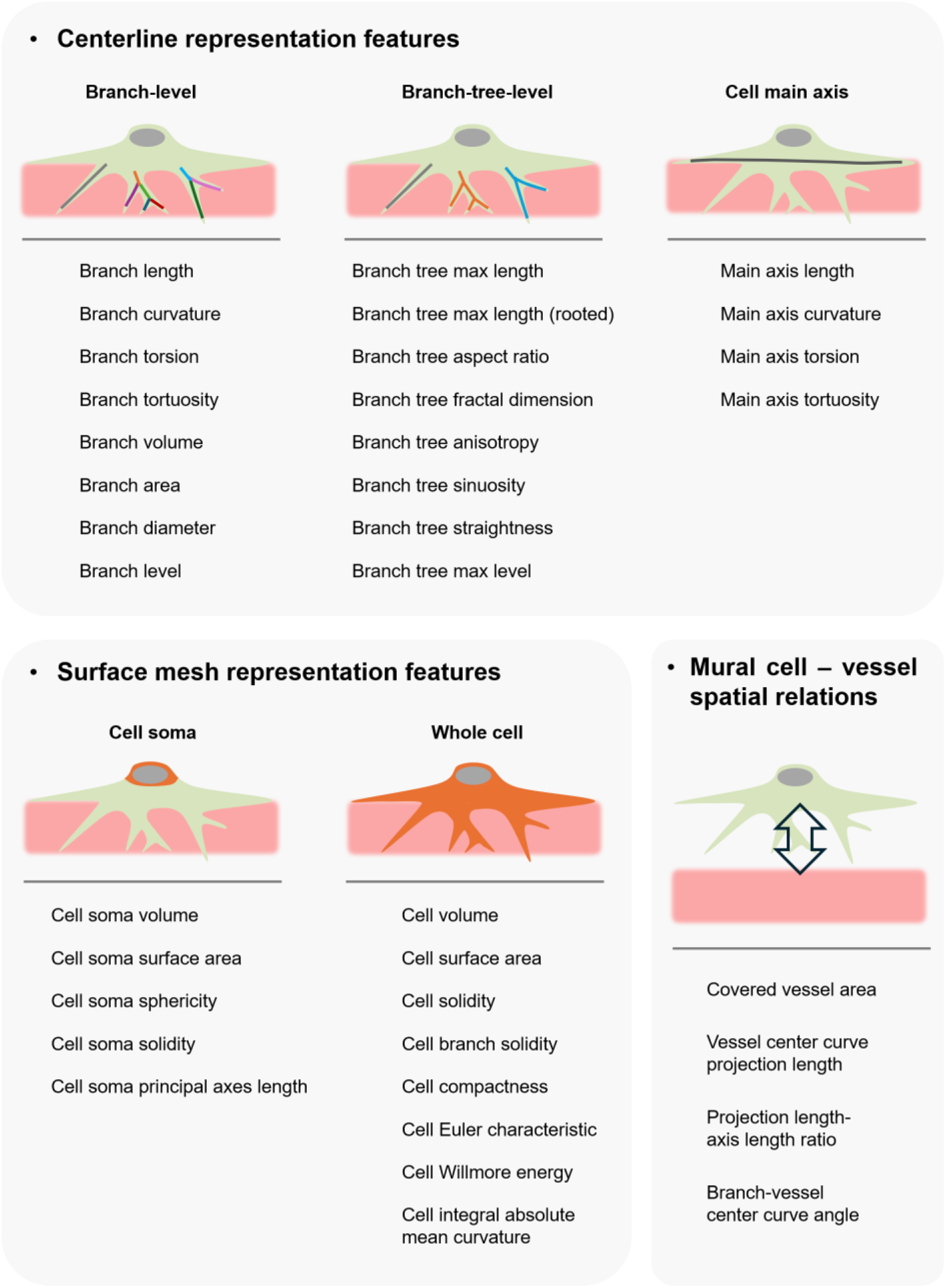
Centerline and surface mesh representation features and mural cell–vessel spatial metrics extracted by Mural-VISTA.

### Zebrafish maintenance and stocks

Zebrafish (*Danio rerio*) were raised and staged according to established protocols [37]. Zebrafish were maintained on a 14-hour light/10-hour dark cycle, and fertilized eggs were collected and raised in E3 medium at 28°C. To inhibit pigmentation in embryos older than 24 hpf, 0.003% N-Phenylthiourea (Sigma-Aldrich, Cat# P7629) in E3 medium was used. *Tg(fli1ep:Lifeact-EGFP)* or *Tg(kdrl:EGFP)* transgenic zebrafish was used to visualize vascular structures. All animal experiments were approved by the Institutional Animal Care and Use Committee at RIKEN Kobe Branch (IACUC).

### Generation of zebrafish Rho CA and Rho DN overexpressing mural cells

To examine how RhoA activity influences cell morphology, a Gal4/UAS system was used to drive RhoA CA or RhoA DN transgene expression transiently and mosaically in mural cells. Plasmid encoding pDestTol2-6xUAS:RhoA G14V-P2A-Lifeact:mCherry (RhoA CA) or pDestTol2-6xUAS:RhoA T19N-P2A-Lifeact:mCherry (RhoA DN) was injected into *TgBAC(pdgfrb:Gal4FF)^ncv24^* zebrafish embryos together with Tol2 transposase mRNA as described in Figure S1. To characterize control mural cells, plasmid encoding pDestTol2-6xUAS: Lifeact:mCherry was used.

### Three-dimensional surface mesh reconstruction

Single mural cell z-stack images were obtained with a confocal microscope in z-directional interval ∼0.30 µm and xy-plane resolution 0.16 µm/pixel setting at 3 days post-fertilization (dpf). Deconvoluted and denoised images were processed using IMARIS (v.9.0.0, BitPlane, Zurich, Switzerland), a 3D image visualization and processing software. Images were binarized according to a built-in, self-adapting algorithm. Triangular surface meshes of vessels and mural cells were created with a 0.50 µm and 0.16 µm surface-detail resolution setting, respectively.

### Comparison with existing analysis methods

Three representative pericyte samples in total from each RhoA condition group were selected as example data. For the cross-method and intra-user test, data was measured by a researcher experienced in biological image data analysis and familiar with all 5 analysis methods (Mural-VISTA, Simple Neurite Tracer (SNT), MotiQ, 3DMorph and Freehand projection image tracing (FPIT)). Three PhD-level researchers with different levels of prior knowledge in bioimage analysis took part in the inter-user test. After learning how to use the compared measurements (Mural-VISTA, SNT and FPIT) on another training image data, examinees were told to process the same three sample images. Reconstructed meshes and .tif files were given as the input. The less experienced researchers received introductory guidance from the experienced researcher and subsequently processed the datasets independently. The time that beginners spent on the training sample and the measurement learning was recorded as the learning time.

### Statistics

Example images of control, RhoA CA-overexpressing, and RhoA DN-overexpressing mural cells were used to illustrate morphological features and the analysis workflow. Trunk mural cells at 3 days post-fertilization (dpf) were used to characterize topo-morphological differences in cell shape resulting from altered RhoA activity (Figure 7 and Figure S2). Control trunk vSMC and pericyte at 3 dpf were used to present the processing steps and outputs of Mural-VISTA program (Figure 4).

**Figure 3.**
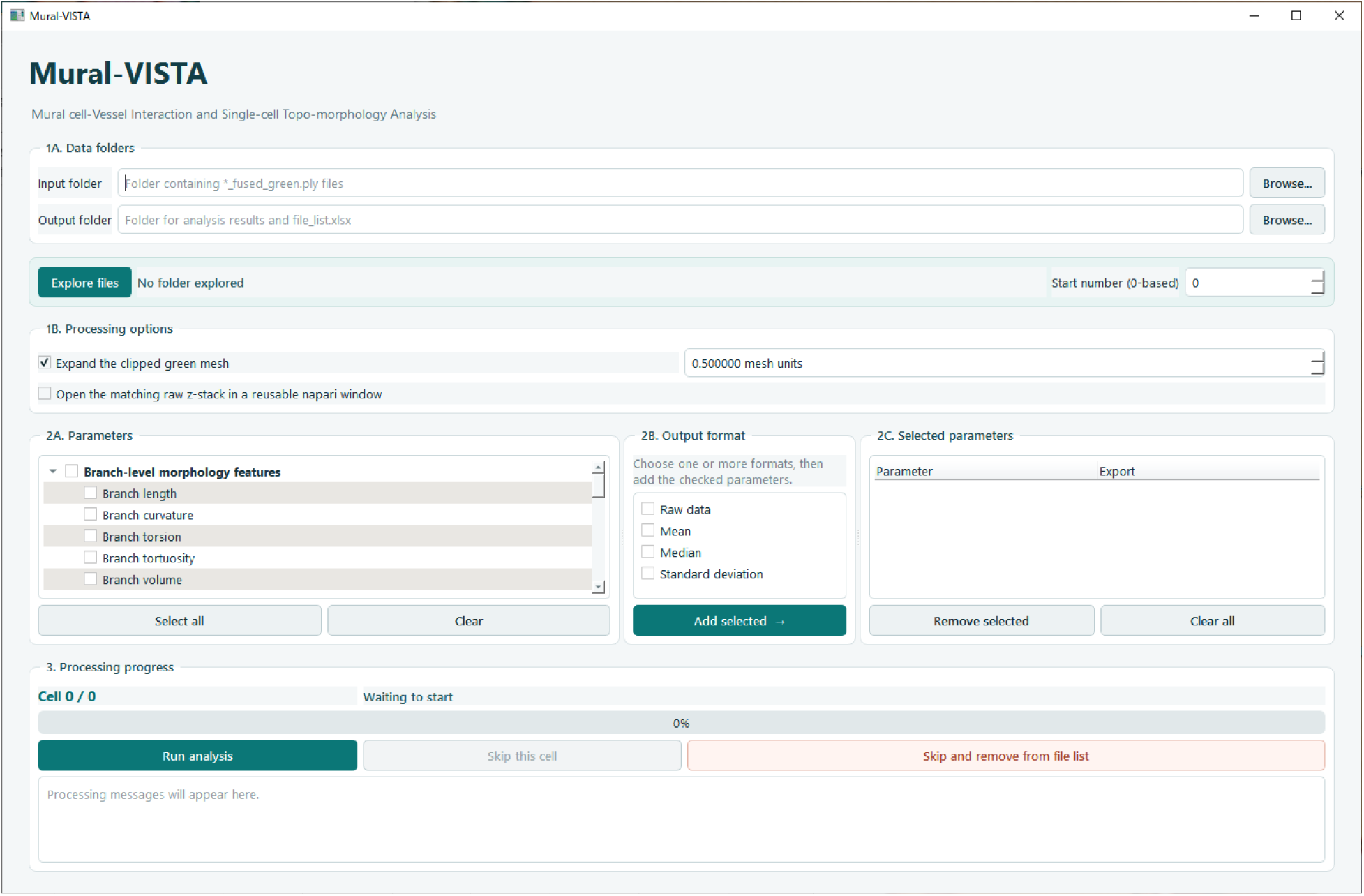
Graphical user interface of the Windows executable file version of Mural-VISTA.

**Figure 4.**
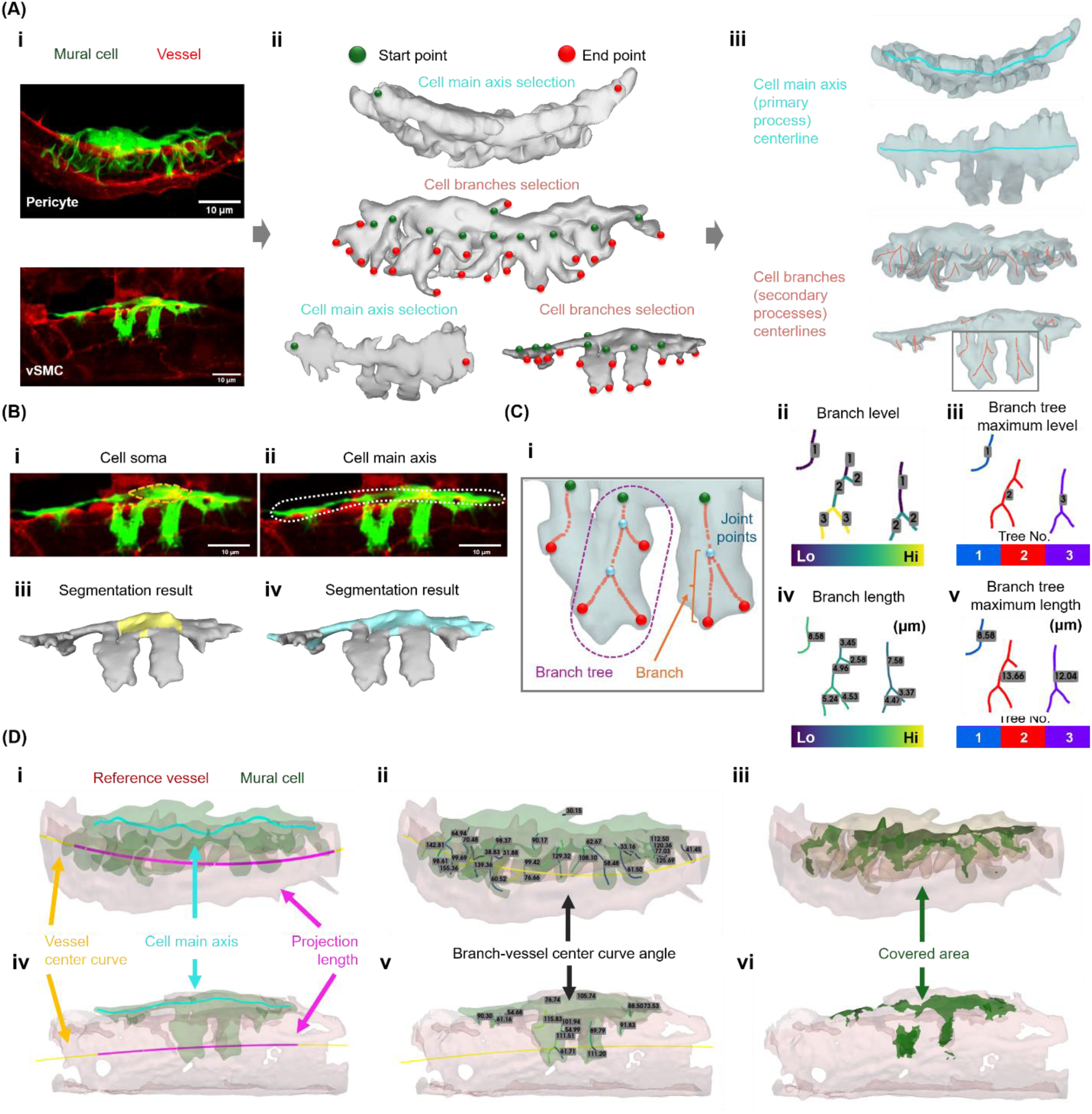
Representative workflow for extracting mural-cell morphological and vessel-referenced spatial features from paired mural-cell and vessel surface meshes. (A) Representative fluorescence image of control pericyte (upper) and vSMC (lower) at 3 dpf (lifeact-mCherry pseudocolored in green) and associated vessel (*fli1ep:Lifeact-EGFP* pseudocolored in red) (i). Screenshots of centerline extraction workflow, from the raw z-stack image (i), reconstructed mesh, start/end point selection (ii) to centerline extraction (iii). (B) Identification of the soma (i) and main axis (ii) in a representative control vSMC and the corresponding segmentation result of soma (iii, yellow) and main axis (iv, cyan). (C) Definition of cell branch structure (i). Representative centerline analysis results of branch level (ii), branch tree maximum level (iii), branch length (iv) and branch tree maximum length (v). (D) Representative vessel-referenced measurements for the pericyte (i–iii) and vSMC (iv–vi): projected whole-cell length along the vessel centerline (i, iv), branch–vessel centerline angle (ii, v), and cell–vessel coverage area (iii, vi).

For quantitative comparisons of trunk pericyte, 26 control cells at 3 dpf from 13 embryos in 3 independent experiments, 32 RhoA CA-overexpression cells at 3dpf from 14 embryos in 2 independent experiments, and 26 RhoA DN-overexpression cells at 3 dpf from 17 embryos in 2 independent experiments were analyzed.

For quantitative comparisons of trunk vSMCs, 28 control cells at 3 dpf from 13 embryos in 3 independent experiments, 38 RhoA CA over-expression cells at 3 dpf from 18 embryos in 2 independent experiments, and 23 RhoA DN over-expression cells at 3 dpf from 13 embryos in 2 independent experiments were analyzed (Figure 8).

Datasets used to establish Mural-VISTA and for quantifying mural cell shape after manipulating RhoA activity are derived from another study [38].

T-tests with Bonferroni correction and one-way repeated-measures ANOVA was used to measure significant differences between analysis methods within each sample–parameter combination. Tukey-Kramer Post hoc test with a one-way ANOVA were used to measure significant differences between mural cells. All statistical differences were considered significant when p-values were < 0.05. Legends used in the figures: *: p< 0.05; **: p< 0.01; ***: p< 0.001; ****: p< 0.0001.

Further details regarding the methods can be found in the supplemental information.

## RESULTS

### Mural-VISTA enables multiscale analysis of mural cell morphology and cell–vessel spatial relationships

We developed Mural-VISTA (Mural cell–Vessel Interaction and Single-cell Topo-morphological Analysis), an interactive Python-based workflow for individual mural cell morphology and mural cell-vessel spatial relationship measurement. This workflow contains five function components : 1) surface mesh loading and trimming, 2) mural cell centerline extraction, 3) mural cell mesh segmentation, 4) multiscale mural cell morphology and mural cell-vessel spatial relation metrics calculation, and 5) result exportation (Figure 1). We also developed a Windows executable file (.exe) version for users without a ready-to-use Python environment (Figure 3).

Mural-VISTA accepts paired surface meshes of a mural cell (compulsory) and its associated vessel (optional), generated from 3D z-stack images by public Python libraries (e.g. Trimesh, VTK/PyVista and PyMeshLab) or commercial software. Users interactively select the start/end points to extract branches (secondary processes) and the cell main axis (primary process) (Figure 4A). The interface supports the selection of multiple endpoints and allows users to inspect and confirm the initial centerline and segmentation results (Figure S4). The extracted centerlines guide segmentation of branches, cell main axis and soma (Figure 4B). Also, they provide the basis for subsequent processing such as segmentation, connectivity searching and branch-level morphological feature measurement mentioned in Figure 2 (Figure 4C). Besides, with the vessel mesh reference, Mural-VISTA calculates the mural cell whole cell projection length and the angles between branch segments and the vessel central curve (Figure 4D-i, ii, iv and v). The coverage position and area on vessel can be calculated as well (Figure 4D-iii and vi). Surface-derived metrics are additionally calculated for the soma and whole-cell mesh. Collectively, the workflow outputs 37 different topological, morphological, and vessel-referenced metrics (Figure 2).

Table 3 summarizes the features of Mural-VISTA in comparison with selected morphology-analysis tools developed for neuronal or microglial analysis and with manual tracing of projected images. Mural-VISTA retains 3D surface representation alongside centerline information in the processing, supports segmentation of the soma, main axis, and branches, and integrates branch-, branch-tree-, soma-, whole-cell-, and vessel-referenced measurements. This combined representation is useful for mural cells, because their processes may include broad or sheet-like regions that are incompletely represented by a centerline alone. The associated vessel mesh further provides a spatial reference for measuring cell orientation, projected length, and vessel coverage. The semi-automatic segmentation therefore combines user-guided definition and confirmation of complex structures with automated metric calculation and result export.

**Table 3.**
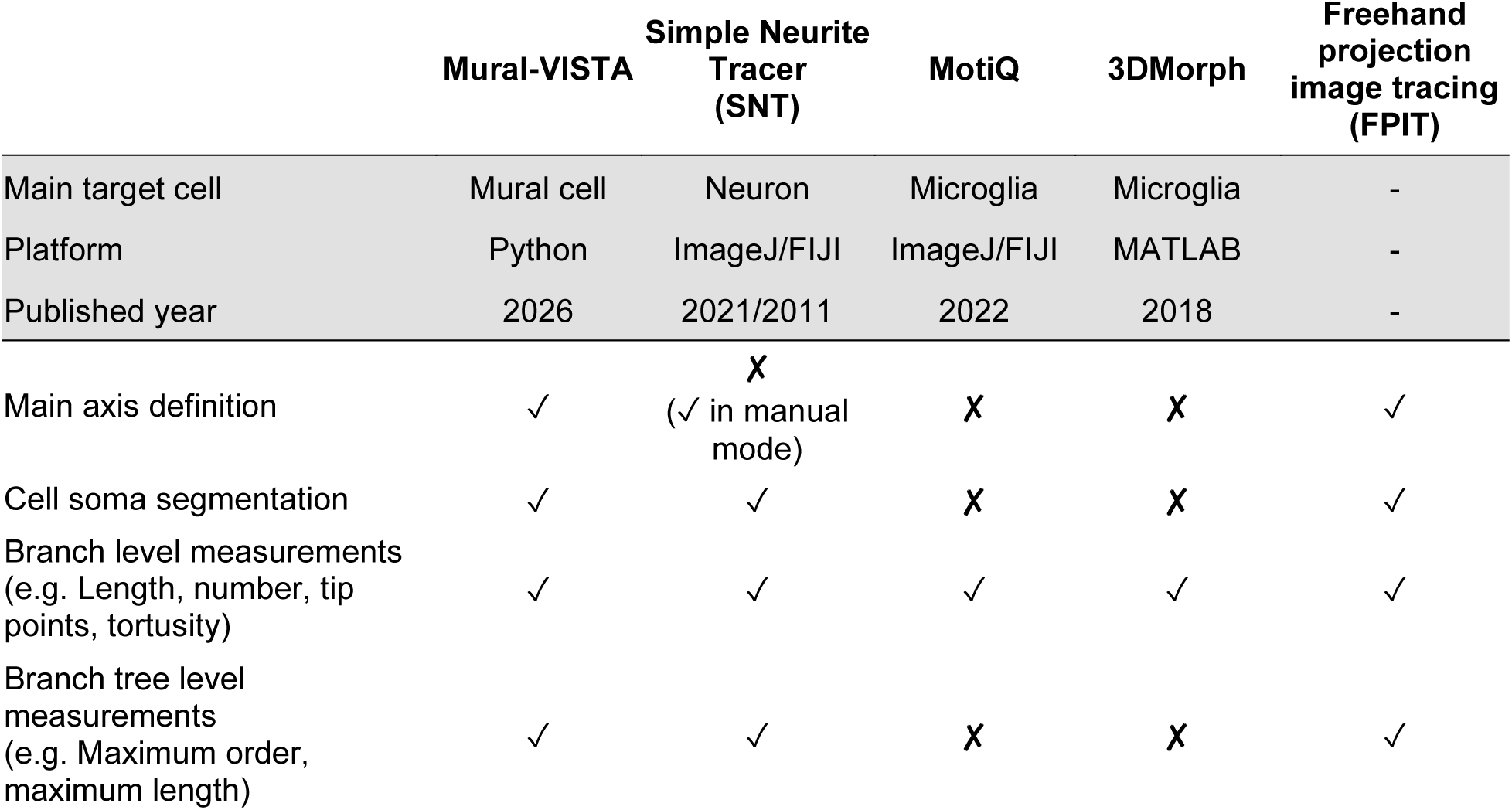
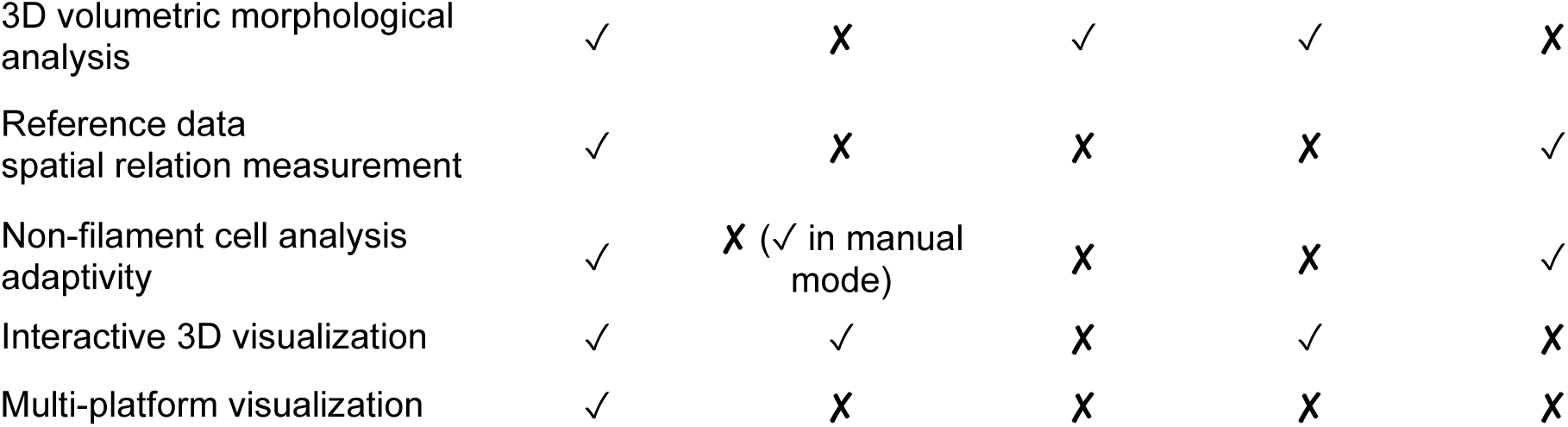
Feature comparisons between Mural-VISTA, selected published analysis tools and freehand manual tracing method.

### Mural-VISTA matches current analysis tools and enables measurement consistency across researchers

We compared Mural-VISTA with existing analysis methods in terms of learning time, processing time, and quantitative measurement outputs. (Figure 5 and 6). Cross-method results showed that Mural-VISTA provides topo-morphological parameters in both centerline and surface mesh representations. Additionally, it provides manually confirmed mesh segmentation for specific regions (cell soma, main axis, and branches) and between branches, enabling further analysis and presentation (Figure 5B). Quantitative analysis showed that Mural-VISTA results matched established methods level and presented the cellular structure (Figure 5C and D).

**Figure 5.**
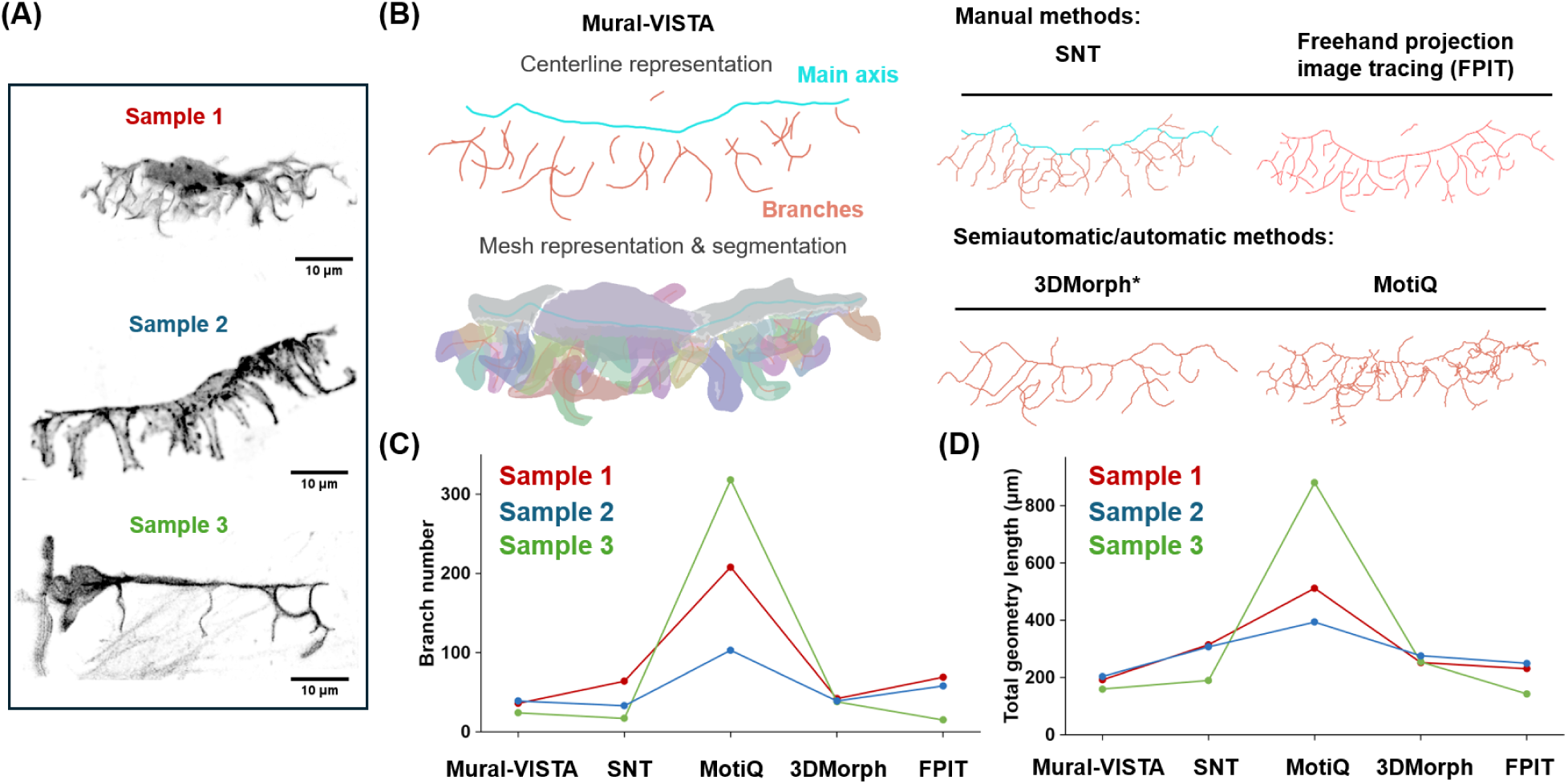
Cross-method comparison between Mural-VISTA and established analysis tools. (A) Maximum intensity projection of the example pericyte z-stack images. (B) Projection of centerline skeletons and mesh segmentation generated by Mural-VISTA and current analysis tools. Main axis (cyan) and branches (red) are separated by different colors in the results that contain hierarchy information. A custom script was used to export skeletons generated by 3DMorph. (C) Branch number comparison between analysis tools’ skeleton results. (D) Total geometry length comparison between analysis tools’ skeleton results.

**Figure 6.**
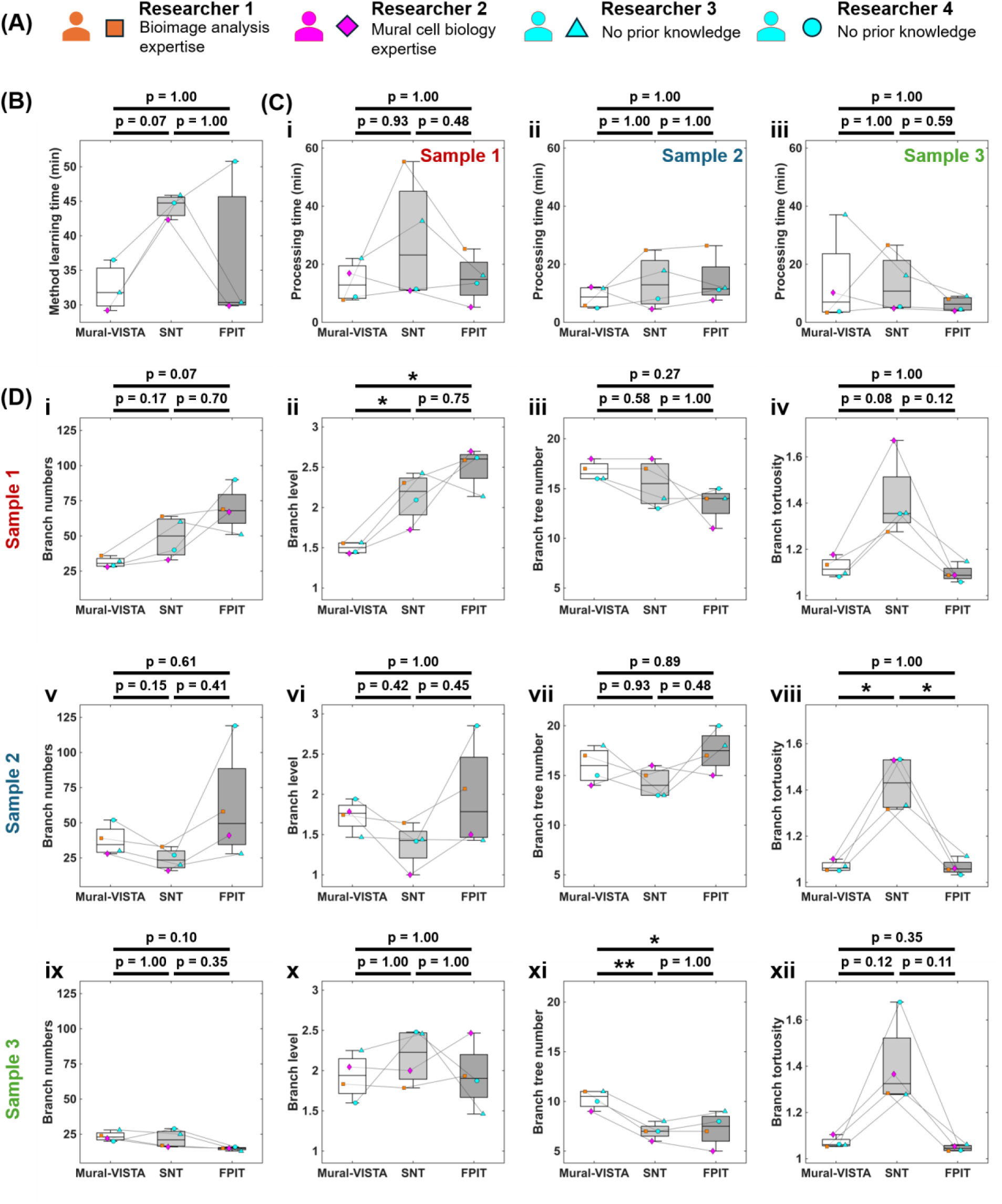
Intra- and inter-user comparison between Mural-VISTA and established analysis tools. (A) Backgrounds of the participating researchers. (B) Learning times among first-time users. (C) Manual processing time between different researchers, method and sample combinations. (D) Topological and morphological metric comparison between tools and researchers, including branch number (i, v, and ix), branch level (ii, vi, and x), branch tree number (iii, vii, and xi) and branch tortuosity (iv, viii, and xii).

For the inter-user test, the same dataset was further analyzed by three independent PhD-level researchers, one with prior knowledge in mural cell biology (data colored magenta) and two without any prior knowledge in biological data processing (data colored cyan). Their results were compared with those of the experienced researcher who did the intra-user test (data colored orange) (Figure 6A). Results showed that the learning times and manual processing time of Mural-VISTA method are almost equal to the conventional methods (Figure 6B and C). However, FPIT outputs required additional post-processing, which was not included in the measured hands-on processing time. Processing times varied across researchers and with sample complexity, but Mural-VISTA narrowed the time variance between experienced people and beginners even in the complex case (Figure 6C-i). The same robustness is also presented in the extracted results. Mural-VISTA results matched those of established analysis tools, with a small inter-user variance. In this study, we applied a 0.5 µm surface offset to all surface mesh, which may contribute to the loss of some topological details (Figure 6D-ii). Regarding the branch sinuosity, we think the surface mesh input benefited the smoothness of the extracted skeleton, while the current method relies on the fluorescence intensity and may need post-treatment (Figure 6D-viii).

### Mural-VISTA resolves cell-type-dependent mural cell morphological responses to RhoA activity

RhoA–ROCK–MLCP–MLC is a canonical pathway where RhoA signaling regulates myosin II phosphorylation level, subsequently manifests in contractile network construction, mural cell morphology and contractility [39,40] (Figure 7A). We have previously observed that control pericytes lining zebrafish intersegmental vessels (ISVs) generate two types of secondary processes – rod-like and lamellar (Figure 7B-i, iv, and vii) [38]. RhoA CA-overexpressing pericyte branches were mainly rod-like, morphologically compact with prominent Lifeact-labeled actin bundles (Figure 7B-iii, vi and ix) [38]. On the contrary, RhoA DN-overexpressing pericyte branches tend to show a lamellar type of shape, morphologically loose and with heterogeneous Lifeact labeling and regions of lower fluorescence intensity (Figure 7B-ii, v and viii) [38]. Given that mural cell secondary process morphology is determined by basal actin filament organization (Figure 7C), and that RhoA activity regulates actin organization and function (Figure 7D), the distinct process morphology may reflect alterations in RhoA activity. We next used Mural-VISTA to determine whether the visually apparent differences were captured by quantitative three-dimensional morphological metrics.

**Figure 7.**
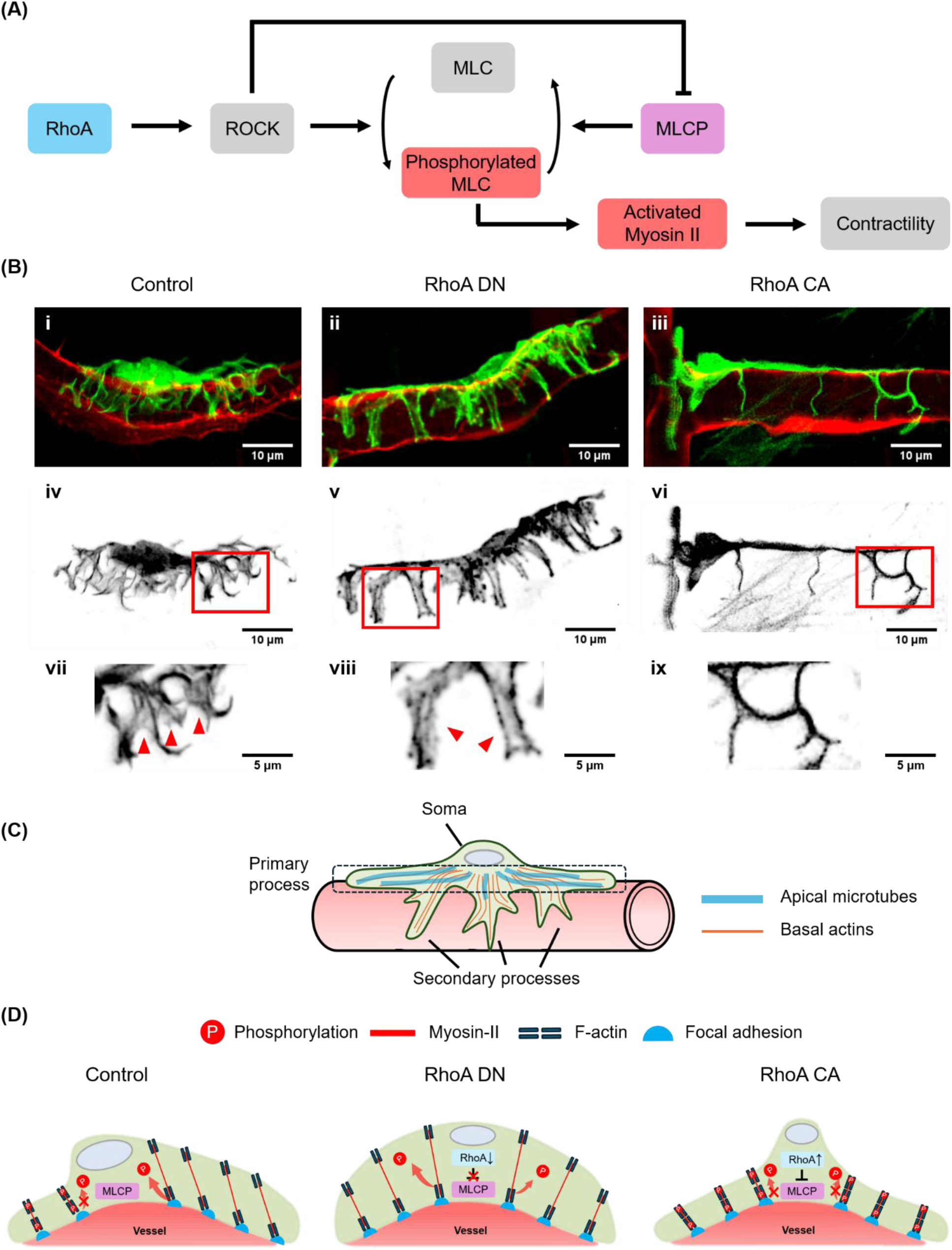
RhoA activity regulates mural cell contractility and morphology. (A) Schematic illustration of the canonical RhoA–ROCK–MLCP–MLC pathway linking RhoA activity to myosin II-dependent contractility. (B) Representative fluorescence image of pericyte (lifeact-mCherry, pseudocolored in green) and associated vessel (fli1ep:Lifeact-EGFP, pseudocolored in red) of control (i), RhoA DN-overexpressing (ii) and RhoA CA-overexpressing pericyte (iii). Grayscale and magnified images of F-actin in control (iv and vii), RhoA DN-overexpressing (v and viii) and RhoA CA-overexpressing pericyte (vi and ix). Pericyte and vessel images were obtained at 3 dpf. Scale bars, 10 µm in i–vi and 5 µm in vii–ix. (C) Schematic illustration of the mural cell hierarchical organization, including the soma, primary process or main axis, and secondary processes. (D) Schematic illustration linking altered RhoA activity to actomyosin organization and secondary-process morphology.

To quantify RhoA-associated morphological phenotypes, we used Mural-VISTA to characterize cell shape, focusing specifically on the complexity of secondary processes. The analysis compared controls with RhoA CA- and RhoA DN-expressing cells and included surface-based solidity metrics, branch abundance and hierarchy, and branch-tree shape metrics (Figure 8). The resulting profiles revealed both shared and cell-type-specific responses to RhoA perturbation.

In pericytes, cell solidity did not differ significantly among the three groups (Figure 8A-i). Cell branch solidity was lower in RhoA DN-overexpressing pericytes than in both controls and RhoA CA-overexpressing pericytes (Figure 8A-ii), consistent with a more spatially dispersed process architecture. RhoA CA overexpression reduced branch number, branch level, and branch-tree maximum level relative to the control (Figure 8B and Figure 8C-i), indicating lower branch abundance and hierarchical complexity. By contrast, RhoA DN overexpression increased branch number but did not significantly alter branch level or branch-tree maximum level relative to the control. Thus, RhoA DN overexpression increased branch abundance without increasing the depth of the branching hierarchy.

**Figure 8.**
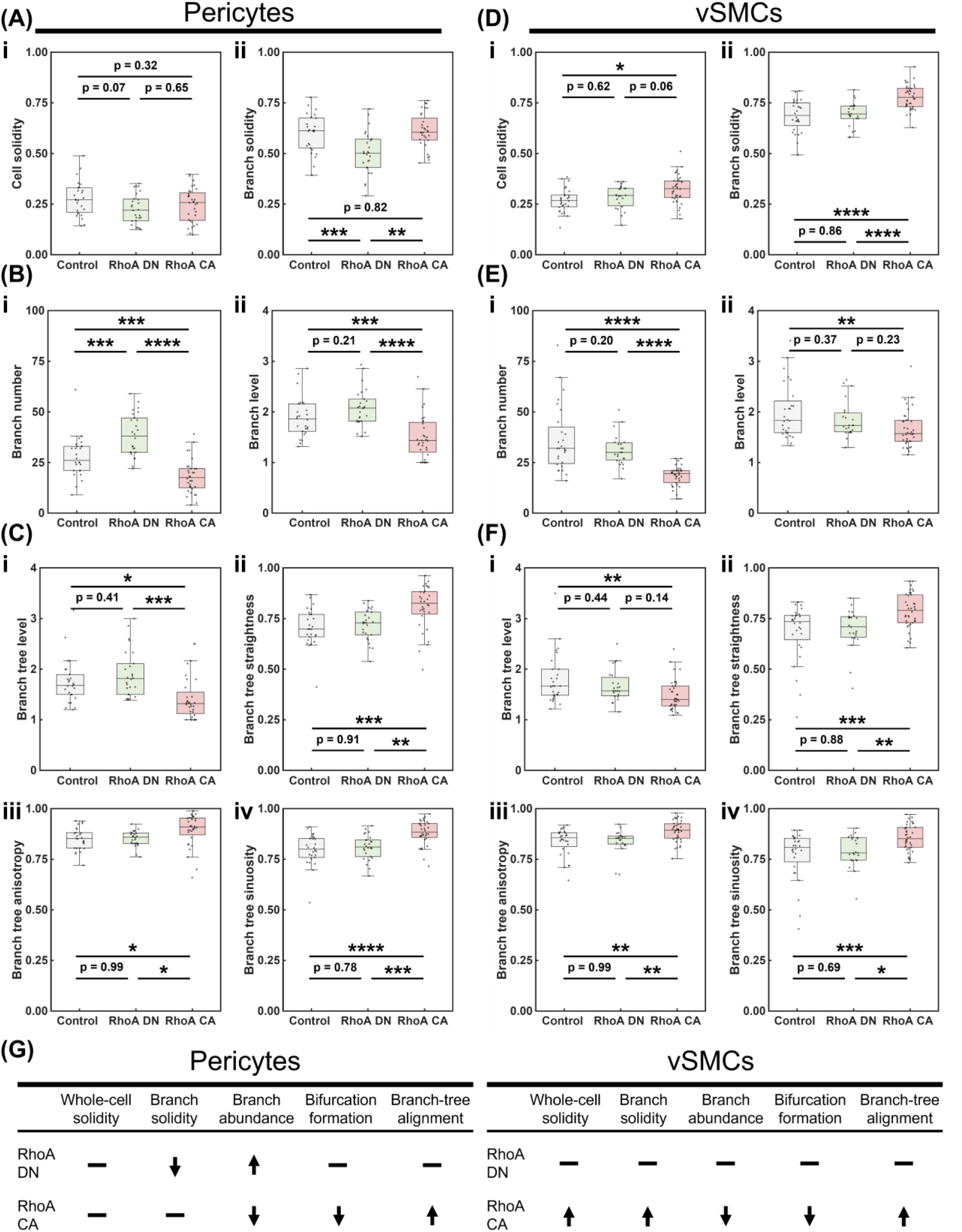
Mural-VISTA reveals mural cell-type-dependent morphological responses to RhoA perturbation. (A) Quantification of surface-mesh-based whole-cell-level solidity metrics, cell solidity (i) and cell branch solidity (ii), of pericytes at 3dpf. (B) Quantification of centerline-based branch-level complexity metrics, branch number (i) and branch level (ii), of pericytes at 3dpf. (C) Quantification of centerline-based branch-tree-level complexity and bendiness metrics, branch tree level (i), branch tree straightness (ii), branch tree anisotropy (iii) and branch tree sinuosity (iv), of pericytes at 3dpf. (D) Quantification of surface-mesh-based whole-cell-level solidity metrics, cell solidity (i) and branch solidity (ii), of vSMCs at 3dpf. (E) Quantification of centerline-based branch-level complexity metrics, branch number (i) and branch level (ii), of vSMCs at 3dpf. (F) Quantification of centerline-based branch-tree-level complexity and bendiness metrics, branch tree level (i), branch tree straightness (ii), branch tree anisotropy (iii) and branch tree sinuosity (iv), of vSMCs at 3dpf. (G) Summary of RhoA CA- and RhoA DN-associated morphological changes in nascent pericyte (i) and vSMC (ii) relative to the control group. Up arrow, down arrow and vertical bar indicate increases, decreases and no significant effect, respectively.

Branch-tree shape metrics further distinguished the pericyte groups. RhoA CA-overexpressing pericytes showed higher branch-tree straightness, anisotropy and sinuosity than controls, indicating stronger global alignment (Figure 8C-ii, iii and iv). RhoA DN-overexpressing pericytes did not differ significantly from controls in branch-tree straightness, anisotropy, or sinuosity. Therefore, the principal RhoA DN-associated pericyte phenotype consisted of increased branch number and reduced branch solidity, consistent with greater process dispersion but not altered branch-tree directional organization. Whereas RhoA CA expression was associated with reduced branching complexity and coordinated changes in branch-tree geometry.

In vSMCs, RhoA CA overexpression increased cell solidity relative to the control and increased cell branch solidity relative to both the control and RhoA DN overexpression conditions (Figure 8D), consistent with a more compact whole-cell and process architecture. RhoA CA also reduced branch number, branch level, and branch-tree maximum level relative to the control (Figure 8E and Figure 8F-i). As in pericytes, RhoA CA-overexpressing vSMCs also showed higher branch-tree straightness, anisotropy, and sinuosity than controls (Figure 8F-ii, iii and iv), indicating processes that were both more globally aligned and locally straighter. None of the measured vSMC metrics differed significantly between the RhoA DN overexpression and control groups.

Together, these results show that constitutively active RhoA reduced branching complexity in both mural cell types, while increasing whole-cell and branch compactness specifically in vSMCs. Dominant-negative RhoA produced a pericyte-selective phenotype characterized by increased branch abundance and lower branch solidity, consistent with greater process dispersion but not altered branch-tree directional alignment. Few detectable changes were observed in vSMCs (Figure 8G).

## DISCUSSION

Here, we developed Mural-VISTA, a semi-automated workflow that integrates 3D surface-mesh analysis, centerline-based topology, hierarchical cell segmentation, and vessel-referenced spatial measurements for individual mural cells. Mural cells present high topological, morphological and functional heterogeneity, making their morphology difficult to represent using conventional two-dimensional projections or cell-centered measurements alone. By analyzing paired meshes of a mural cell and its associated vessel, Mural-VISTA preserves both cellular geometry and vascular context and generates quantitative metrics at branch, branch-tree, main-axis, soma, whole-cell, and cell-vessel levels.

A central feature of Mural-VISTA is the combination of centerline and surface representations. Centerlines are well suited to quantifying branch connectivity, length, hierarchy, curvature and directional organization, as demonstrated by the established neuron and microglia analysis tools [15-18]. Nevertheless, a centerline representation alone doesn’t fully represent broad or lamellar processes and cannot directly provide surface-derived parameters, such as volume, surface area, solidity or contact area. On the other hand, surface meshes retain these morphological features but cannot describe branch topology by themselves. Mural-VISTA uses widely used Python 3D mesh analysis packages, such as VMTK, VTK, PyVista and Trimesh. It combines the complementary information from both representations while using the associated vessel mesh as a reference for projected length, process–vessel angles, and coverage area. As the dataflow chart in Figure 1 shows, this semi-automatic framework is particularly relevant to mural cells because their processes are organized relative to the geometry of the vessel rather than within an unconstrained three-dimensional space. Moreover, intermediate and final results are exported in universal formats and users can easily visualize them via open-source software like MeshLab and ParaView.

The following cross-method and inter-user comparison provided an initial evaluation of Mural-VISTA’s usability and measurement consistency. In this small-scale evaluation, results show that researchers without prior knowledge in cellular biology and biological image analysis can learn its workflow and obtain results broadly comparable to those of an experienced researcher in a short time. However, systematic validation against ground-truth measurement and further assessment of inter-user and inter-lab reproducibility remain necessary.

Previous studies have reported that mural cell processes provide a structural readout of cell status and response to the local vascular context. Direct endothelial contact has been associated with the process formation [41], whereas the formation of process reflects the cell migratory and sensing potential [42]. On the other hand, cytoskeletal gene perturbation, such as CDC42 and SRF, will disrupt the mural cell process and further affect the adjacent vessel, in both morphology and function aspects [43,44]. Our Mural-VISTA program converts structural phenotypes into quantitative variables that can be compared across molecular perturbations, cell types, developmental stages, or vascular environments.

We demonstrated the application of Mural-VISTA by analyzing zebrafish pericytes and vSMCs overexpressing constitutively active or dominant-negative RhoA. RhoA is a key factor that regulates the phosphorylation of MLC and consequently affects actomyosin-generated force [39,40]. Consistent with a role for RhoA in cytoskeletal organization, altered RhoA activity induced morphological changes in mural cells when compared to controls (Figure 7B and S2).

RhoA CA overexpression was associated with reduced branch number, branch level, and branch-tree maximum level in both mural cell types. It also increased branch-tree straightness, anisotropy, and sinuosity relative to control in both pericytes and vSMCs. In addition, elevated RhoA activity increased cell and branch solidity in vSMCs, whereas neither solidity metric differed significantly between RhoA CA-overexpressing and control pericytes. These results indicate that increased RhoA signaling was associated with lower branching complexity in both mural cell types, whereas the increased cell and branch solidity represented a vSMC-specific compactness phenotype.

RhoA DN overexpression produced a different and pericyte-selective phenotype. In pericytes, decreased RhoA activity increased branch number and reduced branch solidity. The increase in branch number did not increase branch level or branch-tree maximum level, suggesting an increase in branch abundance rather than the formation of a bifurcated branch tree. The reduction in branch solidity is consistent with a less compact branch architecture, but it should not be interpreted as reduced branch-tree straightness or altered directional organization. In vSMCs, none of the analyzed metrics differed significantly between the RhoA DN-overexpression and control groups.

## Limitations

In current version (v2.1.0), we admit that tubular branch volume may not fully characterize the process morphology difference, especially in flat lamellar processes. We also acknowledge that the analysis was highly dependent on raw image and reconstructed mesh quality. Future versions should integrate a self-adaptive mesh reconstruction module and an AI-aided high precision pre-segmentation to reduce manual intervention and ensure reproducible results. Besides, inter-user comparison examinee also reported that a progress map with rollback function will be useful. In the long term, we are also willing to generalize this tool to ordinary 3D cell morphology analysis and integrate our non-rigid mesh registration and mesh growth method [45] into this program to realize spatiotemporal morphology analysis of single cells.

## Conclusion

Mural-VISTA provides a semi-automated framework for quantitative three-dimensional analysis of individual mural cells and their spatial relationships with associated vessels. By integrating centerline topology, surface mesh morphology, hierarchical cell segmentation, and vessel-referenced measurements, the workflow captures complementary features of branch organization, cell shape, and cell–vessel geometry. Application to zebrafish mural cells with altered RhoA activity revealed a shared reduced branching complexity and a vSMC-selective increased compactness under RhoA CA-overexpression, whereas a pericyte-selective increase in branch abundance and process dispersion under RhoA DN-overexpression. These findings demonstrate that Mural-VISTA can resolve cell-type-dependent structural phenotypes that are difficult to distinguish using projected images alone. The workflow therefore provides a quantitative basis for future studies of mural cell heterogeneity, molecular perturbation, and cell–vessel remodeling.

## RESOURCE AVAILABILITY

### Lead contact

Further information and requests for resources and reagents should be directed to and will be fulfilled by the lead contact, Yukiko T. Matsunaga.

### Materials availability

This study did not generate new unique reagents.

### Data and code availability

The software and source code used in this study are publicly available as of the date of publication at https://github.com/HedeleZeng/Mural-VISTA.git (GitHub) and 10.5281/zenodo.22124716 (Zenodo).

The version corresponding to this manuscript is archived as release v.2.1.0.

Any additional information required to reanalyze the data reported in this paper is available from the lead contact upon request.

## ACKNOWLEDGMENTS

H.Z. acknowledges the financial support and the fellowship from Japan Science and Technology Agency (JST) Support for Pioneering Research Initiated by the Next Generation (SPRING) (Grant Number JPMJSP2108). L-K.P. was supported by intramural funding from RIKEN BDR and JSPS Grants-in-Aid for Scientific Research grants (22H022624 and 22H05168). M.H. was supported by RIKEN Junior Research Associate Programme and CSC–OU joint scholarship. We appreciate Dr. Weitao Wang and Dr. Qingyu Zhu for participating the inter-user experiment. We also thank Emi Taniguchi and the RIKEN BDR Research Aquarium for zebrafish husbandy support.

## AUTHOR CONTRIBUTIONS

Conceptualization, H.Z., M.H., L-K.P. and Y.T.M.; Methodology, H.Z.; Investigation, H.Z. and M.H.; Data curation, M.H.; Software, H.Z.; Formal analysis, H.Z. and M.H.; Supervision, Y.T.M. and L-K. P.; Resources, Y.T.M. and L-K. P.; Writing—original draft, H.Z.; Writing – review & editing, H.Z., M.H., L-K.P. and Y.T.M.; Funding acquisition, Y.T.M. L-K. P. and H.Z.. All authors have reviewed the manuscript and approved its contents.

## DECLARATION OF GENERATIVE AI AND AI-ASSISTED TECHNOLOGIES IN THE WRITING PROCESS

During the preparation of this work, H.Z. used ChatGPT (Version 4.1 to 6.1) to accelerate the code implementation. After using this tool, H.Z. reviewed and edited the content as needed and takes full responsibility for the content of the publication.

## SUPPLEMENTAL INFORMATION

**Document S1. Figures S1–S4. Video V1.**

## REFERENCES

[1] Siekmann AF, Biology of vascular mural cells. Development (Cambridge, England) 2023;150:dev200271.

[2] Dessalles CA, Babataheri A, Barakat AI, Pericyte mechanics and mechanobiology. J Cell Sci 2021;134:jcs240226.

[3] Zhu S et al., Versatile subtypes of pericytes and their roles in spinal cord injury repair, bone development and repair. Bone Res 2022;10:30.

[4] Grant RI et al., Organizational hierarchy and structural diversity of microvascular pericytes in adult mouse cortex. Journal of cerebral blood flow and metabolism : official journal of the International Society of Cerebral Blood Flow and Metabolism 2019;39:411–425.

[5] Gluck C et al., Distinct signatures of calcium activity in brain mural cells. Elife 2021;10:e70591.

[6] Shanahan CM, Weissberg PL, Smooth muscle cell heterogeneity: patterns of gene expression in vascular smooth muscle cells in vitro and in vivo. Arteriosclerosis, Thrombosis, and Vascular Biology 1998;18:333–338.

[7] Holm A, Heumann T, Augustin HG, Microvascular Mural Cell Organotypic Heterogeneity and Functional Plasticity. Trends Cell Biol 2018;28:302–316.

[8] Wilhelm I et al., Eppur si muove: the dynamic brain pericyte. Fluids and Barriers of the CNS 2025;22:95.

[9] Poduri A et al., Endothelial cells respond to the direction of mechanical stimuli through SMAD signaling to regulate coronary artery size. Development 2017;144:3241–3252.

[10] Singh T, Jacobs KA, Polacheck WJ, Kutys ML, The Notch1 intracellular domain orchestrates mechanotransduction of fluid shear stress. Life Science Alliance 2026;9:e202503599.

[11] O’Hare M, Arboleda-Velasquez JF, Notch Signaling in Vascular Endothelial and Mural Cell Communications. Csh Perspect Med 2022;12:a041159.

[12] Hartmann DA et al., Pericyte structure and distribution in the cerebral cortex revealed by high-resolution imaging of transgenic mice. Neurophotonics 2015;2:041402.

[13] Ornelas S et al., Three-dimensional ultrastructure of the brain pericyte-endothelial interface. Journal of Cerebral Blood Flow & Metabolism 2021;41:2185–2200.

[14] Leonard EV et al., Regenerating vascular mural cells in zebrafish fin blood vessels are not derived from pre-existing mural cells and differentially require Pdgfrb signalling for their development. Development 2022;149:dev199640.

[15] Longair MH, Baker DA, Armstrong JD, Simple Neurite Tracer: open source software for reconstruction, visualization and analysis of neuronal processes. Bioinformatics 2011;27:2453–2454.

[16] Paris I et al., ProMoIJ: A new tool for automatic three-dimensional analysis of microglial process motility. Glia 2018;66:828–845.

[17] Hansen JN et al., MotiQ: an open-source toolbox to quantify the cell motility and morphology of microglia. Mol Biol Cell 2022;33:ar99.

[18] York EM, LeDue JM, Bernier L, MacVicar BA, 3DMorph Automatic Analysis of Microglial Morphology in Three Dimensions from Ex Vivo and In Vivo Imaging. eNeuro 2018;5:ENEURO.0266-18.2018.

[19] Schroeder W, Martin K, Lorensen B, The Visualization Toolkit, An Object-Oriented Approach To 3D Graphics, 4th ed. ed.Kitware; 2006.

[20] Sullivan CB, Kaszynski A, PyVista: 3D plotting and mesh analysis through a streamlined interface for the Visualization Toolkit (VTK). Journal of open source software 2019;4:1450.

[21] Izzo R, Steinman D, Manini S, Antiga L, The Vascular Modeling Toolkit: A Python Library for the Analysis of Tubular Structures in Medical Images. Journal of Open Source Software 2018;3:745.

[22] Attene M, A lightweight approach to repairing digitized polygon meshes. The Visual Computer 2010;26:1393–1406.

[23] Rouvreau V, Alpha complex, GUDHI User and Reference Manual, GUDHI Editorial Board; 2025.

[24] Feng J, Lin W, Chen C, Fractional box-counting approach to fractal dimension estimation, Proceedings of 13th International Conference on Pattern Recognition, vol. 2 IEEE; 1996. pp. 854-858.

[25] Theodorou DN, Suter UW, Shape of unperturbed linear polymers: polypropylene. Macromolecules 1985;18:1206–1214.

[26] Pattnayak PK, Kumar A, Tomar G, Diffusion Dynamics of Star-Shaped Macromolecules in Dilute Solutions. Macromolecules 2024;57:6657–6665.

[27] Dijkstra EW, A note on two problems in connexion with graphs. Numer Math 1959;1:269–271.

[28] Anderson WD, Greenhalgh AD, Takwale A, David S, Vadigepalli R, Novel Influences of IL-10 on CNS Inflammation Revealed by Integrated Analyses of Cytokine Networks and Microglial Morphology. Front Cell Neurosci 2017;11:233.

[29] Özen I et al., Brain pericytes acquire a microglial phenotype after stroke. Acta Neuropathol 2014;128:381–96.

[30] Jiang X et al., Mapping the Plasticity of Morphology, Molecular Properties and Function in Mouse Primary Microglia. Front Cell Neurosci 2022;15:811061.

[31] Davis BM, Salinas-Navarro M, Cordeiro MF, Moons L, De Groef L, Characterizing microglia activation: a spatial statistics approach to maximize information extraction. Sci Rep 2017;7:1576.

[32] Dyn N, Hormann K, Kim S, Levin D, Optimizing 3D triangulations using discrete curvature analysis, Mathematical Methods for Curves and Surfaces: Oslo 2000, Vanderbilt University; 2001. pp. 135 - 146.

[33] Pienaar R, Fischl B, Caviness V, Makris N, Grant PE, A METHODOLOGY FOR ANALYZING CURVATURE IN THE DEVELOPING BRAIN FROM PRETERM TO ADULT. Int J Imaging Syst Technol 2008;18:42–68.

[34] Schamberger B et al., Curvature in Biological Systems: Its Quantification, Emergence, and Implications across the Scales. Adv Mater 2023;35:2206110.

[35] Agus M, Gobbetti E, Pintore G, Cali C, Schneider J, WISH: efficient 3D biological shape classification through Willmore flow and Spherical Harmonics decomposition, 2020 IEEE/CVF Conference on Computer Vision and Pattern Recognition Workshops (CVPRW), IEEE; 2020. pp. 4184-4194.

[36] van Bavel C, Thiels W, Jelier R, Cell shape characterization, alignment, and comparison using FlowShape. Bioinformatics 2023;39:btad383.

[37] Kimmel CB, Ballard WW, Kimmel SR, Ullmann B, Schilling TF, Stages of embryonic development of the zebrafish. Dev Dynam 1995;203:253–310.

[38] Hu M et al., Mural cell contractility regulates vessel diameter by controlling cell morphology and vessel coverage. bioRxiv 2026:2026.06.07.730635.

[39] Nunes KP, Webb RC, New insights into RhoA/Rho-kinase signaling: a key regulator of vascular contraction. Small GTPases 2021;12:458–469.

[40] Somlyo AP, Somlyo AV, Signal transduction and regulation in smooth muscle. Nature 1994;372:231–236.

[41] Brandt MM et al., Transcriptome analysis reveals microvascular endothelial cell-dependent pericyte differentiation. Sci Rep-Uk 2019;9:15586.

[42] Payne LB et al., Pericyte migration and proliferation are tightly synchronized to endothelial cell sprouting dynamics. Integr Biol-Uk 2021;13:31–43.

[43] Orlich MM et al., Mural Cell SRF Controls Pericyte Migration, Vessel Patterning and Blood Flow. Circ Res 2022;131:308–327.

[44] Álvarez-Aznar A et al., Cdc42 is crucial for mural cell migration, proliferation and patterning of the retinal vasculature. Vasc Pharmacol 2025;159:107472.

[45] Zeng H, Sano T, Kawabe J, Matsunaga YT, Spatiotemporal analysis of pericyte-induced nascent angiogenic morphogenesis dynamics and heterogeneity using microvessel-on-a-chips. Cell Biomaterials 2026;2:100216.

