## Supplemental Document S1 for "Mural-VISTA: a tool for mural cell-vessel interaction assessment and multiscale single-cell topo-morphological analysis"

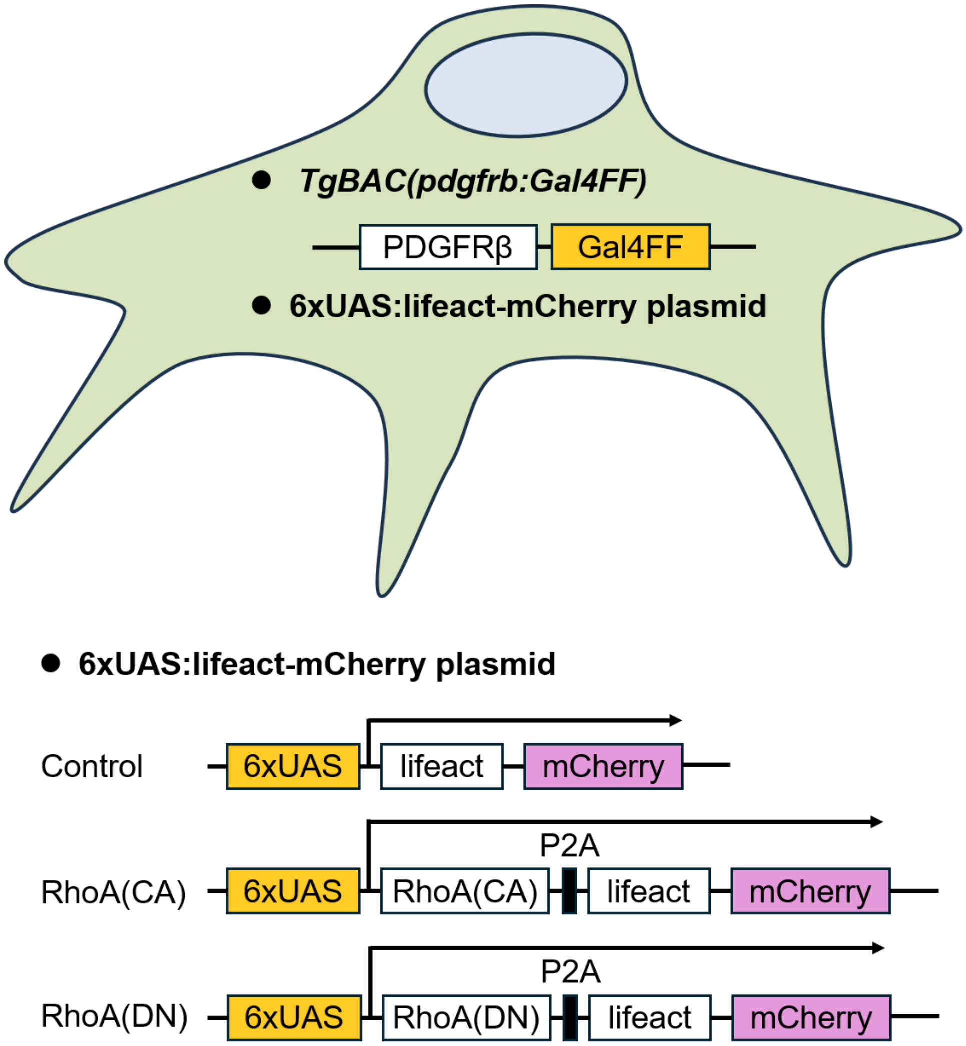


Figure S1. Schematic illustration of the Gal4/UAS setups for mural cell labeling and RhoA level regulation.

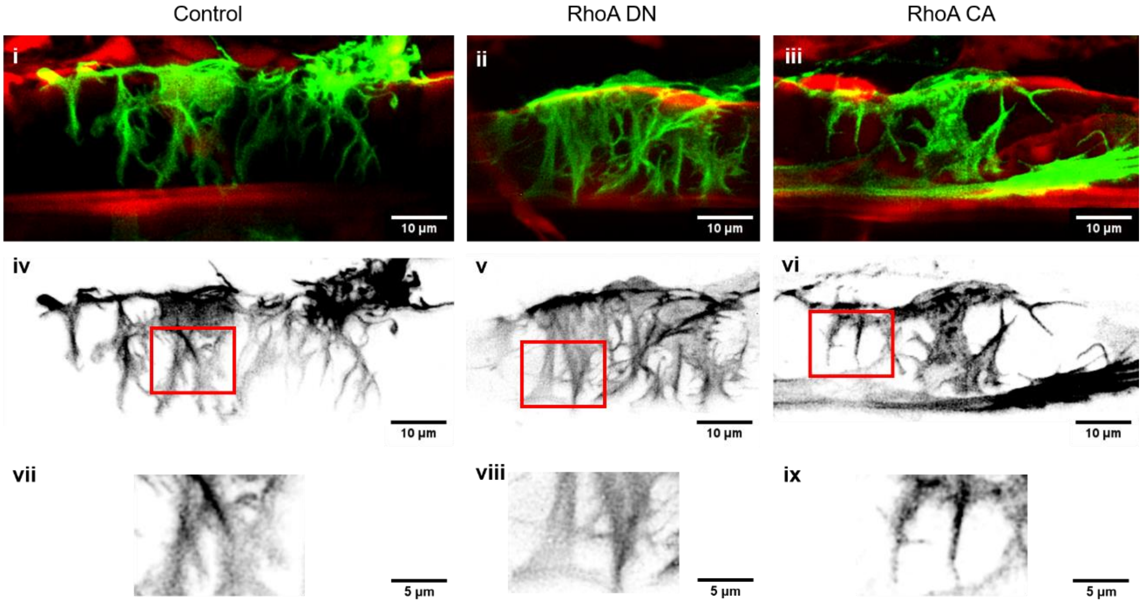


Figure S2. Representative fluorescence image of vSMC (lifeact-mCherry, pseudocolored green) and associated vessel (Fli1ep:Lifeact-EGFP, pseudocolored red) of control (i), RhoA DN-overexpressing (ii), and RhoA CA-overexpressing vSMCs (iii). Grayscale images and zoom-in images of branch F-actin in control (iv and vii), RhoA DN-overexpressing (v and viii) and RhoA CA-overexpressing vSMCs (vi and ix). vSMCs and vessel images were obtained at 3 dpf. Scale bars, 10 µm in i–vi and 5 µm in vii–ix.

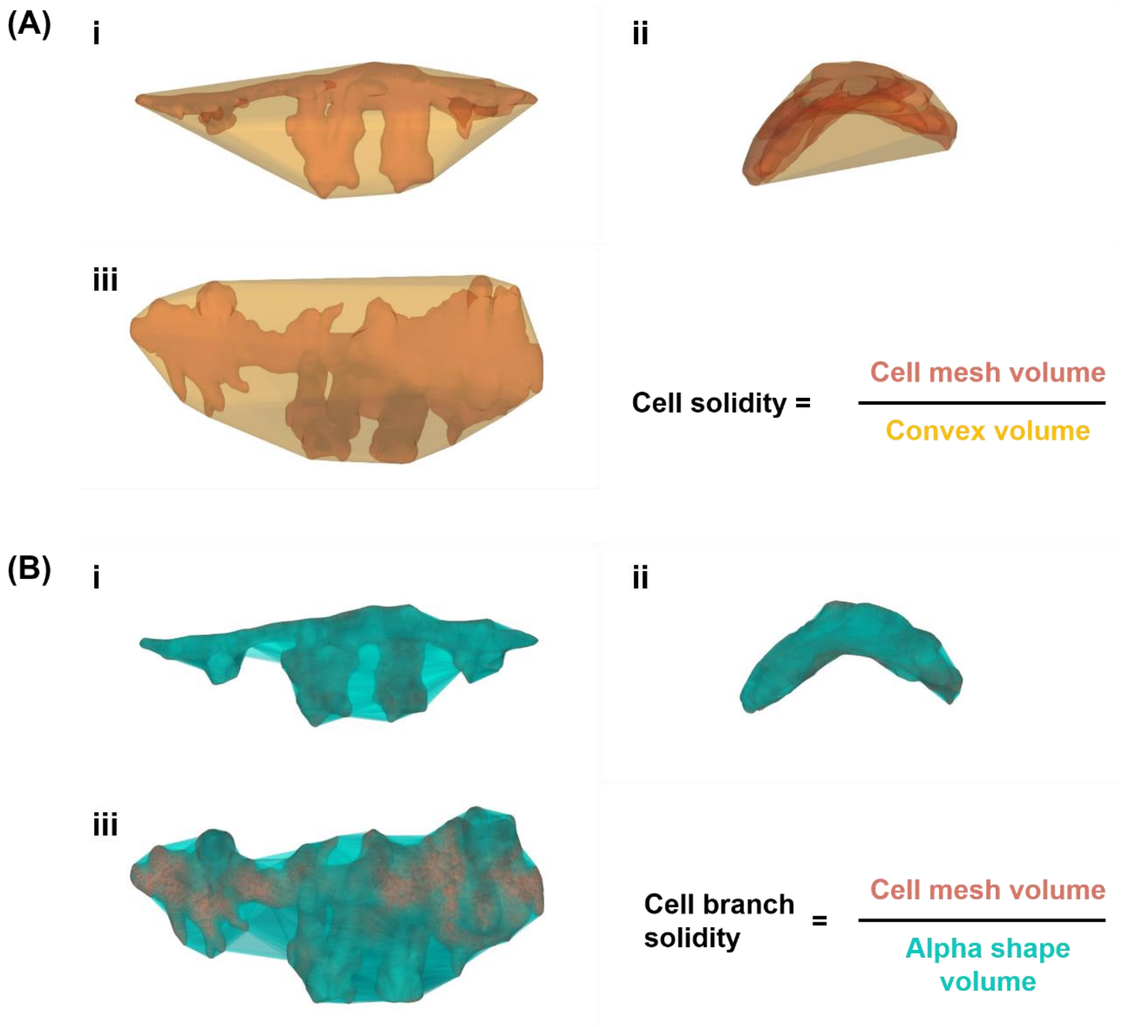


Figure S3. Representative images of cell solidity (A) and cell branch solidity (B) calculation.

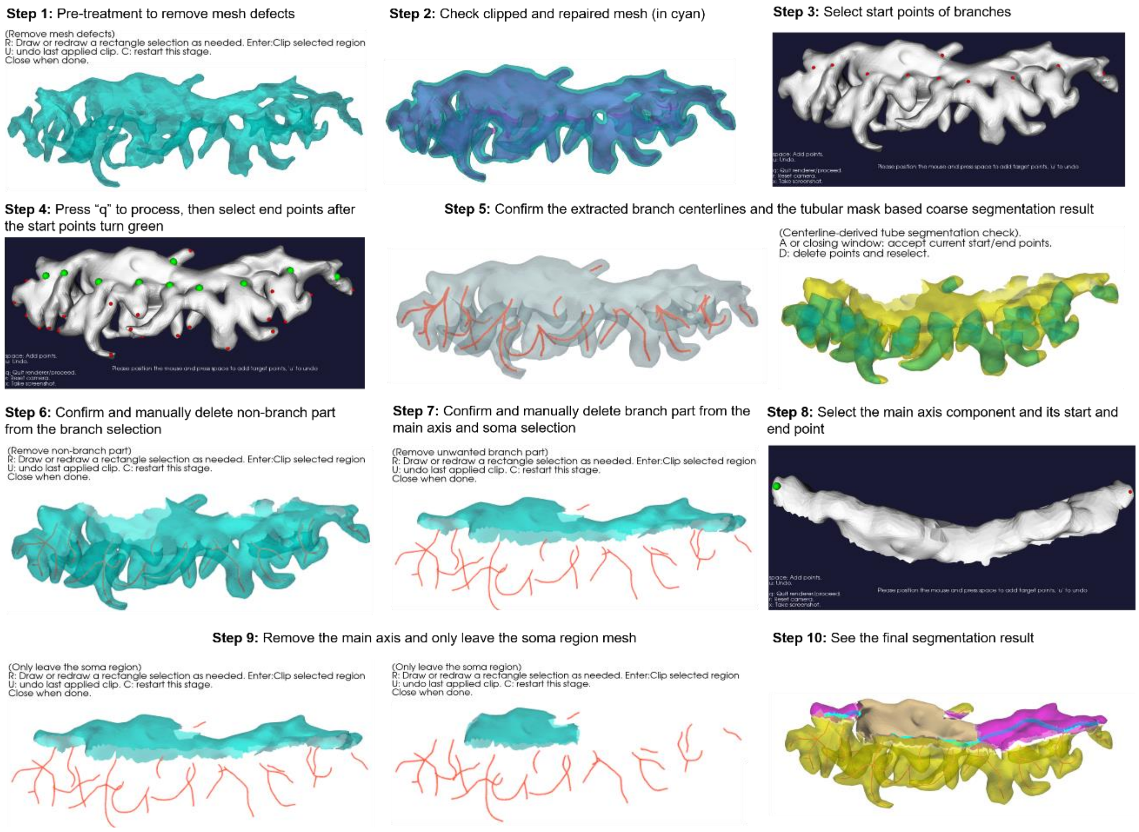


Figure S4. Screenshots and introductions of Mural-VISTA processing steps.
